# Natural abundance isotope ratios to differentiate sources of carbon used during tumor growth in vivo

**DOI:** 10.1101/2020.09.22.307587

**Authors:** Petter Holland, William M. Hagopian, A. Hope Jahren, Tor Erik Rusten

**Affiliations:** Centre for Cancer Cell Reprogramming, Institute of Clinical Medicine, Faculty of Medicine, University of Oslo, Montebello, N-0379 Oslo, Norway; Department of Molecular Cell Biology, Institute for Cancer Research, Oslo University Hospital, Montebello, N-0379 Oslo, Norway; Centre for Earth Evolution and Dynamics, University of Oslo, Blindern, N-0315 Oslo, Norway

**Keywords:** Metabolite, flux, SIRMS, carbon, food, host, tumor, CATSIR

## Abstract

**Background:** Radioactive or stable isotopic labeling of metabolites is a strategy that is routinely used to map the cellular fate of a selected labeled metabolite after it is added to cell culture or to the circulation of an animal. However, the transformation of the labeled metabolite by cellular metabolism within organs complicates the use of this experimental strategy to quantify and understand metabolite transfer between organs. These methods are also technically demanding, expensive and potentially toxic. To allow quantification of the bulk movement of metabolites between organs, we have developed a novel application of stable isotope ratio mass spectrometry (SIRMS).

**Results:** We exploit natural differences in ^13^C/^12^C ratios of plant nutrients for a low-cost and non-toxic carbon labeling, allowing a measurement of bulk carbon transfer between organs *in vivo*. SIRMS measurements were found to be sufficiently sensitive to measure organs from individual *Drosophila melanogaster* larvae, giving robust measurements down to 2.5 µg per sample. We apply the method to determine if carbon incorporated into a growing solid tumor is ultimately derived from food or host tissues.

**Conclusion:** Measuring tumor growth in a *D*.*melanogaster* larvae tumor model reveals that these tumors derive a majority of carbon from host sources. We believe the low cost and non-toxic nature of this methodology gives it broad applicability to study carbon flows between organs also in other animals and for a range of other biological questions.

## Background

An expanding solid tumor has an extraordinary requirement for nutrients to supply anabolic biosynthetic pathways, mostly in the form of carbohydrates and amino acids. The degree to which solid tumor progression depend on nutrients from host feeding is variable for different types of tumors and the metabolic context of the containing organ^1^. Another potential source of nutrients for the tumor is the host itself, obtained by phagocytosis of neighboring cells (entosis), macropinocytosis^2^ or driving release of nutrients from other nearby or distant cells^3^. We set out to determine if carbon incorporated by an expanding tumor is ultimately sourced from ingested food or existing host tissues using a well-established *Drosophila melanogaster* malignant tumor model driven by clonal expression of oncogenic Ras^V12^ and loss of the tumor suppressor scribble.

We sought a method that is agnostic to the identity of incorporated carbon metabolites and the modification of metabolites in metabolic pathways of different organs *in vivo*. Existing methods like radioactive ^14^C-tracing or ^13^C detection through mass spectrometry could allow us to follow a selected metabolite either by feeding or infusing it and then looking for the label in the tumor, but existing applications of these methods do not allow a measurement of carbon transfer from host tissues to the tumor. Moreover, labeled metabolites are expensive and only allow relative measurements between samples for one metabolite per experiment, not absolute measurements of the mass transfer of carbon.

To allow this type of measurement we developed an experimental methodology that we have named CArbon Transfer measured by Stable Isotope Ratios (CATSIR), which exploits differences in the abundance of ^13^C/^12^C of biomolecules in edible plants to allow low-cost and non-toxic tracking of the carbon in metabolites. The ^13^C/^12^C is expressed in the delta notation (*δ*^13^C) in units of per mille (‰) and reported relative to the international standard Vienna Peedee Belemnite (VPDB). Plants can be categorized into two main groups with distinct variants of photosynthesis that results in different *δ*^13^C values of the resulting plant biomolecules. The two groups are called C3-type with *δ*^13^C ≅ -25‰ (i.e potato and beets) and C4-type with *δ*^13^C ≅ -12‰ (i.e. corn and sugar cane)^4^. The absolute differences in ^13^C between C3 and C4 plants are small, but can be accurately quantified by stable isotope ratio mass spectrometry (SIRMS), routinely performed by biologists and biogeochemists to study these plants and how they interact with their environment.

## Results

*D*.*melanogaster* larvae require a food source containing carbohydrates, amino acids and lipids for optimal growth. This is achieved by mixing sources of sugar, complex carbohydrates and yeast with agar to create a pellet of food where eggs are laid and the larvae develop. We determined the commercial baker’s yeast that we use in our standard fly food as being similar to other C3-type nutrients and used this as a C3-type yeast, mixed with potato mash and beet sucrose to create C3 food (Figure 1 a). To generate C4 yeast we expanded commercial yeast on sugar cane sucrose as the carbon source and mixed the C4 yeast with sugar cane sucrose and corn flour to create C4 food. By having flies lay eggs on this food and allowing the larvae to develop, we obtained fully C3- or C4-labeled larvae (Figure 1 b). The lower limit of carbon required to obtain reliable *δ*^13^C measurements was found to be around 2.5 micrograms of carbon per sample, allowing us to reliably measure organs from individual animals. The tumor measurements are performed by extracting the cephalic complex containing the brain as well as the eye discs where the tumor is growing in our genetic tumor model. It is necessary to take the whole cephalic complex because as the tumor expands, it outgrows the eye discs and invades the brain, making dissection of only the tumor or eye discs impossible at later stages of tumor development.

**Figure 1:**
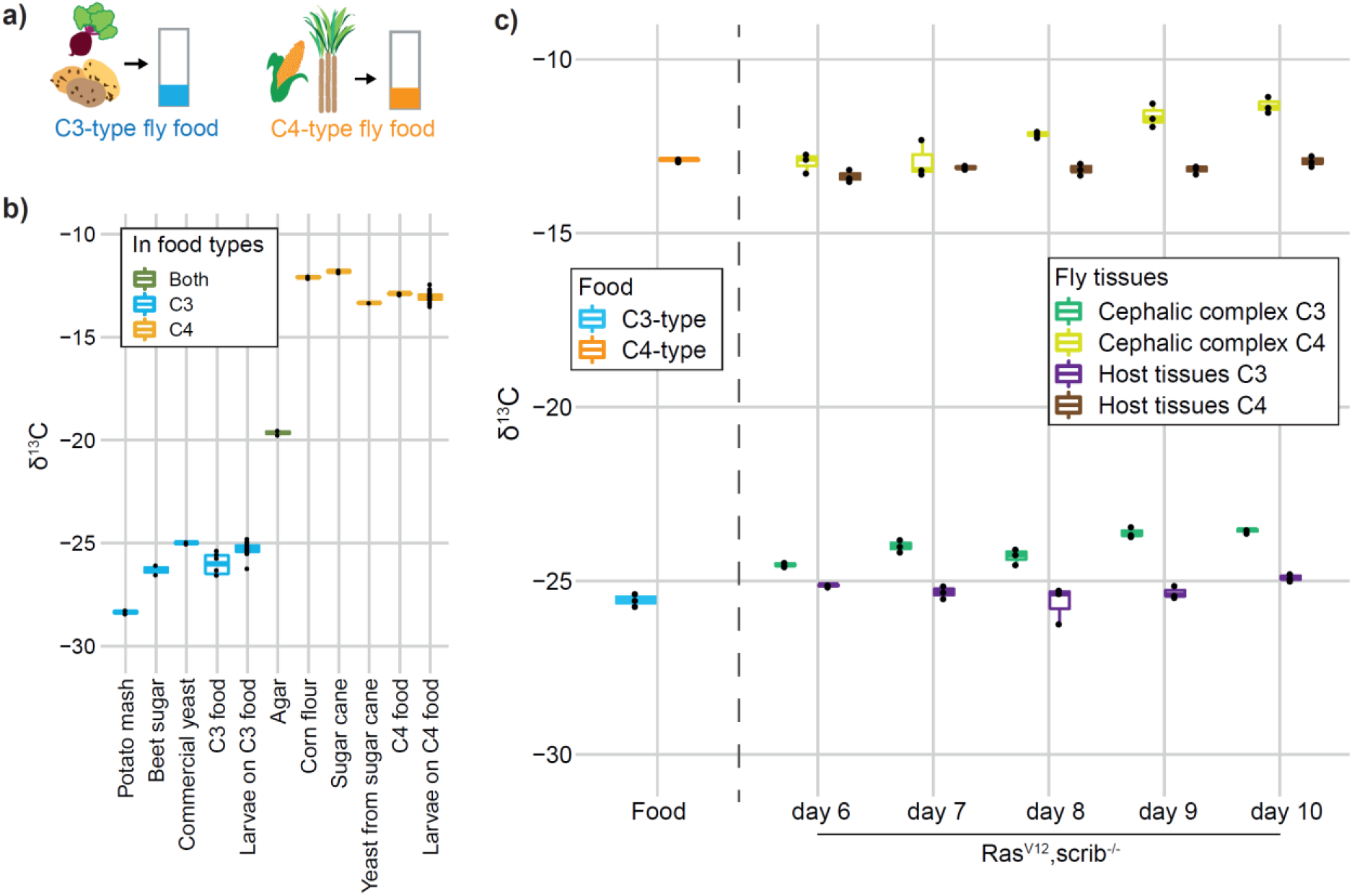
C3- and C4-plant based fly food for stable isotopic labeling of *D*.*melanogaster*. Food composition (a) and *δ*^13^C of the food components and flies developed on the indicated food types (b). Individual food components in b) are measured in duplicate or singlicate (for C4 yeast). The C3 and C4 composite food types are measured with six replicates and the larvae growing on the two food types are measured with 23 biological replicates. c) ^13^C/^12^C measurements of the cephalic complex (eye disc and brain) from fly larvae with genetically programmed (Ras^V12^,scrib^-/-^) tumors in the eye disc. Quantified as *δ*^13^C units per mille (‰), relative to the international ^13^C/^12^C standard Vienna Peedee Belemnite (VPDB). The tumors develop from a transformed group of cells contained within the eye disc at day 6 of larval development to overgrowing the eye disc and invading the brain at day 10. “Host tissues” is a measurement of the remaining organs of the larvae after removing the cephalic complex. Measurements of the food used in these experiments is also included. The larvae organ measurements in c) are from three biological replicates for each indicated time of larvae development. Each datapoint represents the SIRMS measurement of a single animal. Box-plot are used for visualizing the data with default settings for geom_boxplot in R; the median as a line inside boxes extending from the 25^th^ percentile to the 75^th^ percentile and whiskers extend maximally to 1.5x of the inter-quartile range.

In our early trials measuring tumors in animals growing only on either C3 or C4 food we found that as a tumor grows on one type of food, the measured *δ*^13^C of the cephalic complex gradually becomes less negative, while the other host tissues of the same larvae do not change (Figure 1 c). We found the rate and relative amount of ^13^C enrichment by the tumors to be similar for larvae growing on either the C3 or C4 food. A recent study that measured *δ*^13^C of human breast cancer biopsies also found an enrichment of ^13^C in human tumor biopsies relative to neighboring control tissue from the same patient^5^. They observed an enrichment resulting in the *δ*^13^C values to be increased by ∼3‰ compared to adjacent tissue from the same patient, a similar effect size as what we see in our fly model (Figure 1 c). The enrichment of ^13^C by transformed cells was also seen in cell culture of commonly used cell lines of both human and mouse origin^5^ and thus appears to be an inherent feature of transformed cells, but the metabolic reason for tumor ^13^C enrichment remains unknown.

To differentiate if carbon incorporated into an expanding tumor biomass is ultimately derived from ingested food or the host, we need an experimental situation where the carbon in the food is labeled differently than the carbon in host tissues. We can achieve this in the Ras^V12^,scrib^-/-^ *D*.*melanogaster* tumor model if the food source is changed at day 6 (from C3-type to C4-type), when the host tissues are fully developed, but the tumor is very small (Figure 2 a). Pupation (the transformation from larvae to adult fly) normally starts around day 6 for these animals, but the growing tumor delays this process, giving an experimental window of several days starting from day 6 when the food and host tissues will have a different carbon composition, the host tissues are isotopically stable (Figure 1 c), and the tumor is growing exponentially. We illustrate how the *δ*^13^C measurements can be informative to determine the source of carbon used for tissue growth with simulated data in Figure 2 b.

**Figure 2:**
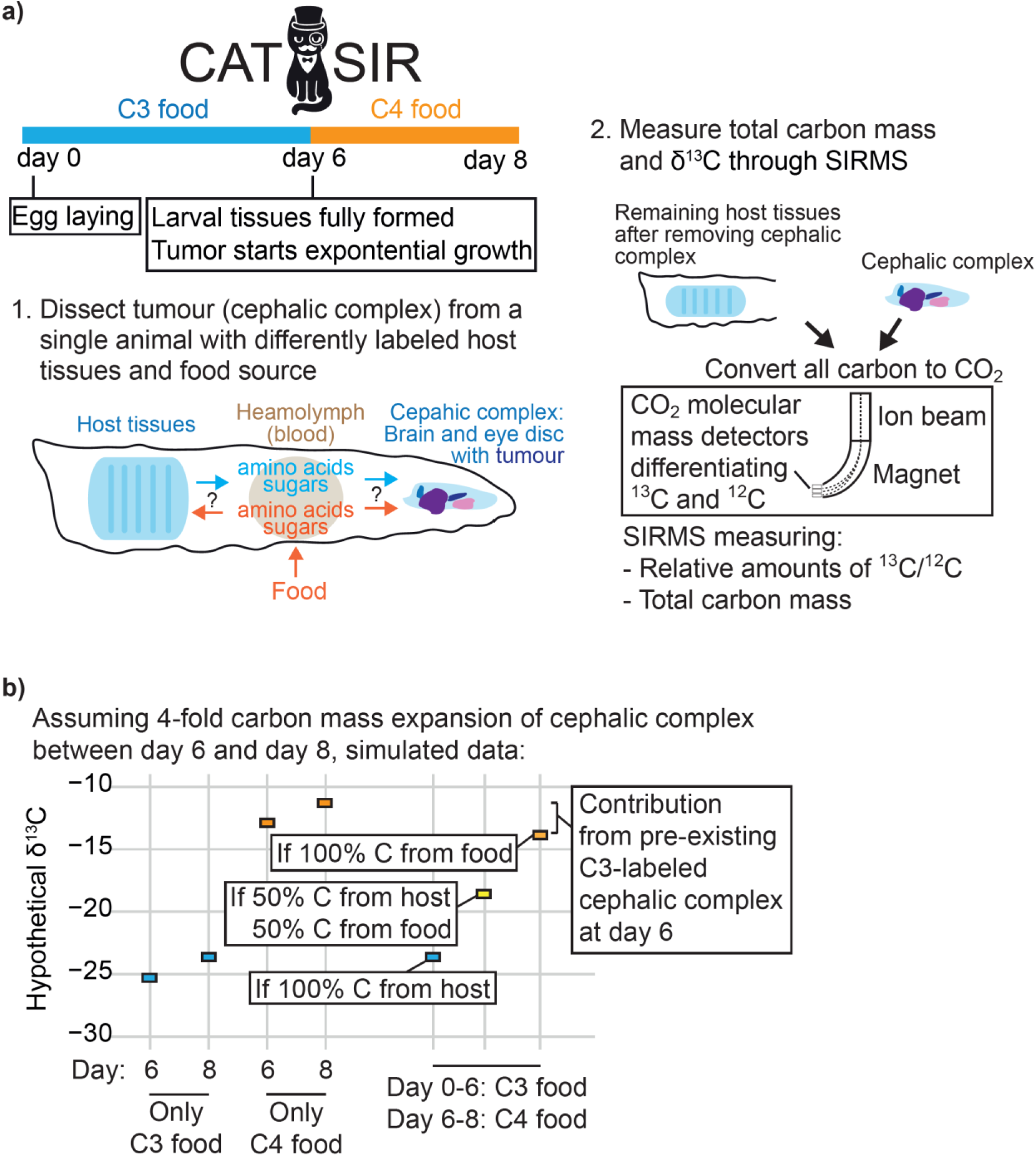
CATSIR, a method to differentiate if carbon incorporated into a growing tumor is derived from ingested food or from host tissues. a) Overview of the experimental setup of the CATSIR methodology. b) Simulated data to illustrate how the *δ*^13^C measurements are used to differentiate the sources of carbon. ^13^C/^12^C measurements are quantified as *δ*^13^C units per mille (‰), relative to the international ^13^C/^12^C standard Vienna Peedee Belemnite (VPDB).

As a consequence of the observed tumor ^13^C enrichment (Figure 1 c), *δ*^13^C measurements of the food itself cannot be used as a basis to calculate how much carbon from ingested food is being incorporated into the growing tumor. Instead, we rely on measuring tumors from animals developing on either of the two food types at multiple stages of development, creating baseline measurements that allow calculations of where a tumor is sourcing its carbon (Figure 3 a). Importantly, we found that the cephalic complex mass is not significantly different between larvae growing only on C3 or C4 food at neither day 6 or 8 (Figure 3 b, p=0.21 at day 6, p=0.21 at day 8), meaning the two food types are similarly able to support tumor growth.

**Figure 3:**
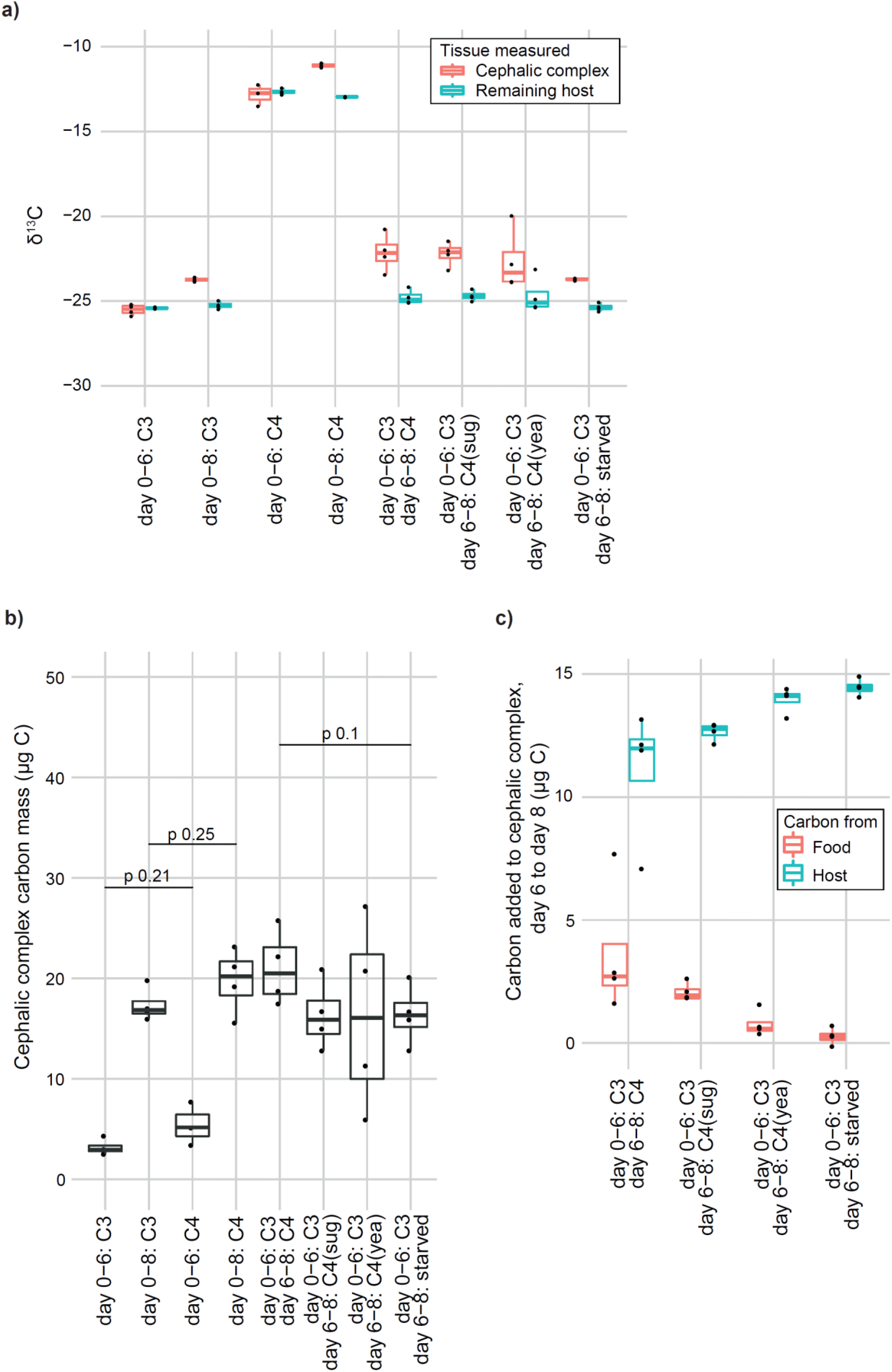
CATSIR applied to measure the sources of carbon used for tumor growth in *D*.*melanogaster* larvae. a) *δ*13C measurements of indicated tissue from Ras^V12^,scrib^-/-^ larvae. In addition to the standard C4 food, larvae were moved to other food variants – one with only sugars (C4 sug), one with only yeast (C4 yea) and one without any nutrients (starved). b) Cephalic complex total carbon mass measurements of the same larvae as in a). c) The calculated amounts of carbon from the food or host tissues being incorporated into the tumor between day 6 and 8, calculated from the SIRMS data shown in a) and b). Measurements in this figure are from four independent biological replicates with the exception of three replicates for the C4 day 6 set where one replicate was lost because there was not enough carbon in it to obtain a reliable *δ*^13^C value. The indicated statistical tests in b) are performed by an unpaired two-sided t-test between the indicated groups. Box-plots are used for visualizing the data with default settings for geom_boxplot in R; the median as a line inside boxes extending from the 25^th^ percentile to the 75^th^ percentile and whiskers extending maximally to 1.5x of the inter-quartile range.

Incorporating the baseline measurements, we then solve two separate equations to calculate where the carbon incorporated into the tumor from day 6 to day 8 is coming from; one determining the isotopic composition of the cephalic complex if the growth was incorporating only food-derived carbon and the other equation gives the isotopic composition of the cephalic complex if the tumor was incorporating only host-derived carbon. For the larvae that was moved from C3 to C4 food on day 6, the experimentally measured isotopic composition of the cephalic complex at day 8 is compared to these two theoretical values to create a factor of the relative contribution from the food and host. The measured carbon mass added between day 6 and day 8 (Figure 3 b) is multiplied by this factor to calculate the mass of carbon added to the tumor from the two sources (Figure 3 c). Using this approach, we found that Ras^V12^,scrib^-/-^ driven tumor growth between day 6 and day 8, tumor growth that causes a 4-fold increase in the cephalic complex carbon mass (Figure 3 b), sources a majority of the carbon it uses for biomass expansion from host sources as well as a smaller amount directly from the food (Figure 3 c).

This methodology can further be adapted to allow measurements of how the different carbon sources in the food change the amount of food-derived carbon that is incorporated in tumor growth. Moving the larvae at day 6 to food with C4 sugar (no yeast) demonstrated a relative contribution to growth from the food and host carbon that is similar to what is seen for a complete C4 food, while having only C4 yeast (no sugars) showed less carbon incorporation into the tumor from the food (Figure 3 c). Having no nutrients (only agar, starved) in the food demonstrated no incorporation of carbon from the food (Figure 3 c). We were surprised to see that the tumor growth itself was not significantly reduced when various nutritional components are removed from the food, even when there are no nutrients (Figure 3 b, p=0.1). This observation, seen together with the large amount of incorporation of host-derived carbon also when food is present, is a key insight from these experiments that point to a close interaction between tumor and host metabolism *in vivo*.

## Discussion

The CATSIR methodology allows a new type of measurement of bulk carbon transfer *in vivo* and expands the methodological arsenal of researchers studying systemic metabolism. Rather than competing with or replacing existing methods, we think CATSIR has strong synergy with other types of metabolite flux measurements that rely on labeling selected metabolites by radioactivity or stable isotopes. CATSIR also uniquely allows a direct measurement of carbon mass, information that is not typically available through detection of specifically labeled metabolites, where the read-out is a relative measurement of the label between samples. We imagine an experimental that strategy that starts with many low-cost CATSIR experiments and then following up on selected candidates with detailed studies using isotopic or radioactive tracers is a way to maximize the biological insights about metabolite flows through systemic metabolism.

The main limiation on the use cases for the CATSIR methodology is that is requires organ growth in the experimental interval when the food is changed. However, the general principle of using C3- and C4-based food to label tissues in vivo and SIRMS measurements allow other types of studies that are not limited to measuring organ growth. One possibility is to mix C3 and C4 food components together and measure the incorporation of carbon from the C3 and C4 sources into different organs, allowing insights about the preference of different organs for different categories of metabolites. Another application is to use the differences in C3 and C4 labeling to give a type of feeding measurement for an organism that goes beyond measuring food ingestion and instead measures incorporation of the food-derived nutrients.

## Conclusion

Here, we succesfully employ CATSIR to measure the mass of carbon incorporated into a growing tumor from host and food sources in a *D*.*melanogaster* tumor model. The demonstrated strategy should be adaptable to in vivo studies of any animal because of the sensitivity, simplicity, low cost and non-toxic nature of the carbon labeling. Through technical optimizations we achieved reliable measurements down to 2.5 µg of total carbon per sample, making the methodology applicable for smaller samples like biopsies that are relatively easily obtained. More generally, the use of C3- and C4-based food to label tissues and subsequent SIRMS measurements is low-cost unexplored experimental strategy that should allow new types of measurements for a range of biological questions.

## Methods

### Fly food

C3 food was prepared with 32.7 g/L potato mash, 60 g/L beet-derived sucrose and 27.3 g/L commercial dry yeast (Lesaffre, Saf-instant. *δ*^13^*C* measured to be similar to C3-type plants). C4 food was prepared with 32.7 g/L corn flour, 62 g/L cane-derived sucrose and 26.3 g/L commercial yeast that was expanded on sucrose from sugar cane. The amount of sugar cane and yeast added to the C4 food was slightly adjusted compared to the C3 food to account for the higher protein content and lower carbohydrate content of the corn flour compared to the potato mash, giving a similar final fat, protein and carbohydrate content of the two food variants. Both foods were also added 4.55 ml/L propionic acid (Sigma, P5561), 2 g/L nipagin (Sigma, H5501) and 7.3 g/L agar (AS Pals, 77000).

### Fly genetics

Larvae with genetically programmed tumors in the eye disc were generated by crossing y,w,ey-flp; Act>y+>Gal4, UAS-GFP/CyO; Frt82B, tub-Gal80 females with y,w;UAS-Ras^V12^/CyO; Frt82B,scrib^-^ /TM6B males.

### Sample preparation

When moving larvae from C3 to C4 food, holes were poked in surface of the new food to give easy immediate access to the new food. Before dissection, larvae were washed 3 times in water and the cephalic complex (eye disc and brain) was dissected in a drop of ultrapure water. The cephalic complex from single larvae was added to a tin capsule (Elemental microanalysis, D1006) and the remaining tissues of the larvae after removing the cephalic complex was added to a separate tin capsule. Samples were then left to dry in a desiccator.

### Stable isotope measurement

The carbon stable isotope value of each sample was determined using a Delta V Advantage Isotope Ratio Mass Spectrometer (Thermo Fisher, Bremen, Germany) configured with a Thermo Fisher EA Isolink Elemental Analyzer at the University of Oslo, Norway. Samples were loaded into a zero blank autosampler (Costech Analytical, Valencia, USA) and quantitatively combusted to CO_2_ via Dumas combustion in the elemental analyzer. The CO_2_ flowed to the mass spectrometer within a stream of helium where the ^13^C/^12^C were determined. Carbon stable isotope values were expressed in the delta notation (δ^13^C) in units of per mille (‰).

Multiple replicates of two internal lab reference materials (“JRICE”, a white rice obtained from a supermarket and homogenized with a ball mill, δ^13^C = -27.43‰ ; and “JGLUT”, L-glutamic acid obtained from Fisher Scientific, δ^13^C = -13.43‰) were incorporated into each analytical batch run and used to normalize the data to the Vienna Pee Dee Belemnite (VPDB) scale using a standard regression method^6^. Additionally, a quality control sample (“JALA”, L-Alanine from Fisher Scientific, δ^13^C = -20.62‰) was incorporated into every batch run and analyzed as an unknown. All three materials (JRICE, JGLUT, JALA) were calibrated within our laboratory and normalized to VPDB using LSVEC and NBS-19, which define the VPDB scale^7^. To verify that our calibrations were accurate, we analyzed IAEA-601 benzoic acid (consensus δ^13^C = -28.81‰) as an unknown and obtained δ^13^C = - 28.83 ± 0.04‰ (1σ., n = 6). Over the course of all sample analyses, the JALA quality control sample returned a mean value of -20.61 ± 0.10 (1σ, n = 45), which is in agreement with our calibrated value of -20.62‰.

We calculated micrograms of carbon in each sample using the Thermal Conductivity Detector (TCD) peak areas from the Flash EA Isolink elemental analyzer. A standard curve was generated from a size series of ACS grade glucose (Thermo Scientific), created using a serial dilution of the glucose in water. Ten microliter aliquots of each solution were added to an empty tin capsule, resulting in a series of capsules containing between 1 and 100 µg of carbon once the water evaporated. This enabled us to determine the amount of carbon in each combusted capsule down to 1.0 µg ± 5%.

### Carbon transfer calculations

To calculate the relative contribution of carbon from the food and host tissues to tumour growth between days 6 and 8, the carbon mass and δ^13^C needs to be measured for larvae growing only on C3 food at day 6 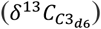 and day 8 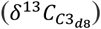 as well as larvae growing only on C4 food at day 8 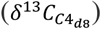. When an experimental larvae is moved from C3 to C4 food at day 6 and then measured at day 8, the amount of tumor growth between day 6 and day 8 is calculated by subtracting the measured cephalic complex carbon mass at day 8 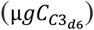 from the mean cephalic complex mass at day 6 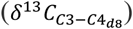 giving μ*gC*_*growth*_. Two theoretical δ^13^C values are then calculated to determine what the isotopic composition would be at day 8 if all the carbon for growth between day 6 and day 8 was coming from the food (δ^13^*C*_*allFood*_) or if all carbon was coming from the host (δ^13^*C*_*allHost*_):

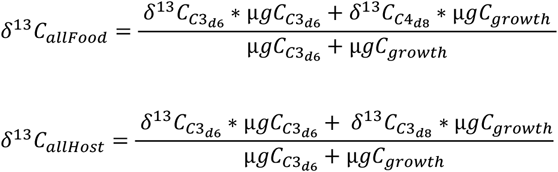

The measured δ^13^C of the cephalic complex at day 8 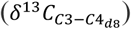 is then compared to these two theoretical values:

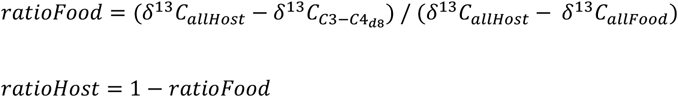

Finally, these ratios are multiplied by the measured carbon mass added to each sample between day 6 and day 8 to derive the total amount of carbon added from the food or host:

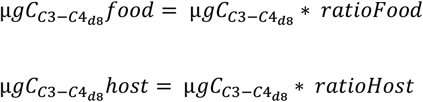

## Supporting information

SIRMS raw data and R scripts for calculations

## Declarations

### Ethics approval and consent to participate

The experiments with Drosophila melanogaster were performed in accordance with The Norwegian Department of Health’s guidelines and approvals for transgenic animals and facilities.

## Availability of data and materials

Raw data from SIRMS measurements as well as R scripts that contain all required calculations and scripts to directly reproduce the figures in the manuscript from the raw data are supplied in the supplementary data.

## Competing interests

The authors declare that they have no competing interests.

## Notes

### Competing Interest Statement

The authors have declared no competing interest.

## References

1. Lien, E. C. & Vander Heiden, M. G. A framework for examining how diet impacts tumour metabolism. Nature Reviews Cancer 19, 651–661 (2019).

2. Palm, W. & Thompson, C. B. Nutrient acquisition strategies of mammalian cells. Nature 546, 234–242 (2017).

3. Katheder, N. S. et al. Microenvironmental autophagy promotes tumour growth. Nature 541, 417–420 (2017).

4. Chartrand, M. M. G. & Mester, Z. Carbon isotope measurements of foods containing sugar: A survey. Food Chem. 300, 125106 (2019).

5. Tea, I. et al. 13C and 15N natural isotope abundance reflects breast cancer cell metabolism. Sci. Rep. 6, (2016).

6. Skrzypek, G. Normalization procedures and reference material selection in stable HCNOS isotope analyses: An overview. Analytical and Bioanalytical Chemistry 405, 2815–2823 (2013).

7. Coplen, T. B. et al. After two decades a second anchor for the VPDB δ13C scale. Rapid Communications in Mass Spectrometry 20, 3165–3166 (2006).

