## Supplementary material for "Natural abundance isotope ratios to differentiate sources of carbon used during tumor growth in vivo": SIRMS raw data and R scripts for calculations: SIRMS calculation R scripts.R

```
options(stringsAsFactors=FALSE)
library(ggplot2)
library(cowplot)
library(readxl)
library(ggsignif)

isDat = list()
isDat[["190425"]] = as.data.frame(read_excel("D:/rad/190425 isotope foods/Larvae Data 051319_PH.xlsx",sheet="Sheet2",skip=0))
isDat[["190521"]] = as.data.frame(read_excel("D:/rad/190521 isotope foods switching/Larvae Data 061419_edited.xlsx",sheet="Sheet1",skip=2))
isDat[["190521"]][,1] = gsub("r","R",isDat[["190521"]][,1])
isDat[["190521"]][,1] = gsub("tb","TB",isDat[["190521"]][,1])
isDat[["190521"]][,1] = sapply(strsplit(isDat[["190521"]][,1]," "),function(x) paste(x,collapse="_"))
colnames(isDat[["190521"]]) = c("ID","d13C","µgC","Notes")
isDat[["190619"]] = as.data.frame(read_excel("D:/rad/190619 isotope food switching 2/Larvae Data 091619_processed.xlsx",sheet="Sheet1"))
isDat[["190619"]][,1] = sapply(strsplit(isDat[["190619"]][,1]," "),function(x) paste(paste(x[seq(1,(length(x)-1))],collapse="_"),x[length(x)],collapse=" "))
colnames(isDat[["190619"]]) = c("ID","d13C","µgC")
isDat[["191031"]] = as.data.frame(read_excel("D:/rad/191031 isotope food switching 3/Larvae Data 170320.xlsx",sheet="Sheet1",skip=21))
isDat[["191031"]][,1] = sapply(strsplit(isDat[["191031"]][,1]," "),function(x) paste(paste(x[seq(1,(length(x)-1))],collapse="_"),x[length(x)],collapse=" "))
isDat[["191031"]] = isDat[["191031"]][,c(1,2,4,5)]
isDat[["191031"]][,1] = gsub("Tb","TB",isDat[["191031"]][,1])
colnames(isDat[["191031"]]) = c("ID","d13C","µgC","Notes")
isDat[["200420"]] = as.data.frame(read_excel("D:/rad/200420 isotope RsTP locRes/Larvae Data 250520.xlsx",sheet="Sheet1",skip=21))
isDat[["200420"]] = isDat[["200420"]][,c(1,2,4,5)]
colnames(isDat[["200420"]]) = c("ID","d13C","µgC","Notes")

# EXCLUDED
isDat[["191031"]] = isDat[["191031"]][isDat[["191031"]][,"ID"]!="R12s_C4_d6_TB 3",] # Below reliable measurement limit


# Food and animal measurements
foodDf = data.frame()
foodDf = rbind(foodDf,cbind("Potato mash",isDat[["190425"]][grep("Potatomash",isDat[["190425"]][,1]),"d13C"],"C3"))
foodDf = rbind(foodDf,cbind("Potato mash",isDat[["190619"]][grep("Potato Potato",isDat[["190619"]][,1]),"d13C"],"C3"))
foodDf = rbind(foodDf,cbind("Beet sugar",isDat[["190425"]][grep("Farin sugar",isDat[["190425"]][,1]),"d13C"],"C3"))
foodDf = rbind(foodDf,cbind("Beet sugar",isDat[["190619"]][grep("Farin Farin",isDat[["190619"]][,1]),"d13C"],"C3"))
foodDf = rbind(foodDf,cbind("Commercial yeast",isDat[["190425"]][grep("Yeast",isDat[["190425"]][,1]),"d13C"],"C3"))
foodDf = rbind(foodDf,cbind("Commercial yeast",isDat[["190619"]][grep("C3 yeast",isDat[["190619"]][,1]),"d13C"],"C3"))
foodDf = rbind(foodDf,cbind("C3 food",isDat[["190619"]][grep("C3_food",isDat[["190619"]][,1]),"d13C"],"C3"))
foodDf = rbind(foodDf,cbind("C3 food",isDat[["200420"]][grep("C3_food",isDat[["200420"]][,1]),"d13C"],"C3"))
foodDf = rbind(foodDf,cbind("Larvae on C3 food",isDat[["191031"]][grep("R12s_C3_d6_R",isDat[["191031"]][,1]),"d13C"],"C3"))
foodDf = rbind(foodDf,cbind("Larvae on C3 food",isDat[["191031"]][grep("R12s_C3_d8_R",isDat[["191031"]][,1]),"d13C"],"C3"))
foodDf = rbind(foodDf,cbind("Larvae on C3 food",isDat[["200420"]][grep("R12s_C3_d6_R",isDat[["200420"]][,1]),"d13C"],"C3"))
foodDf = rbind(foodDf,cbind("Larvae on C3 food",isDat[["200420"]][grep("R12s_C3_d7_R",isDat[["200420"]][,1]),"d13C"],"C3"))
foodDf = rbind(foodDf,cbind("Larvae on C3 food",isDat[["200420"]][grep("R12s_C3_d8_R",isDat[["200420"]][,1]),"d13C"],"C3"))
foodDf = rbind(foodDf,cbind("Larvae on C3 food",isDat[["200420"]][grep("R12s_C3_d9_R",isDat[["200420"]][,1]),"d13C"],"C3"))
foodDf = rbind(foodDf,cbind("Larvae on C3 food",isDat[["200420"]][grep("R12s_C3_d10_R",isDat[["200420"]][,1]),"d13C"],"C3"))
foodDf = rbind(foodDf,cbind("Agar",isDat[["190425"]][grep("Agar",isDat[["190425"]][,1]),"d13C"],"Both"))
foodDf = rbind(foodDf,cbind("Agar",isDat[["190619"]][grep("Agar Agar",isDat[["190619"]][,1]),"d13C"],"Both"))
foodDf = rbind(foodDf,cbind("Corn flour",isDat[["190425"]][grep("Corn flour",isDat[["190425"]][,1]),"d13C"],"C4"))
foodDf = rbind(foodDf,cbind("Corn flour",isDat[["190619"]][grep("Corn flour",isDat[["190619"]][,1]),"d13C"],"C4"))
foodDf = rbind(foodDf,cbind("Sugar cane",isDat[["190425"]][grep("Cane sugar",isDat[["190425"]][,1]),"d13C"],"C4"))
foodDf = rbind(foodDf,cbind("Sugar cane",isDat[["190619"]][grep("Sug cane",isDat[["190619"]][,1]),"d13C"],"C4"))
foodDf = rbind(foodDf,cbind("Yeast from sugar cane",isDat[["190619"]][grep("C4 yeast",isDat[["190619"]][,1]),"d13C"],"C4"))
foodDf = rbind(foodDf,cbind("C4 food",isDat[["190619"]][grep("C4_food",isDat[["190619"]][,1]),"d13C"],"C4"))
foodDf = rbind(foodDf,cbind("C4 food",isDat[["200420"]][grep("C4_food",isDat[["200420"]][,1]),"d13C"],"C4"))
foodDf = rbind(foodDf,cbind("Larvae on C4 food",isDat[["191031"]][grep("R12s_C4_d6_R",isDat[["191031"]][,1]),"d13C"],"C4"))
foodDf = rbind(foodDf,cbind("Larvae on C4 food",isDat[["191031"]][grep("R12s_C4_d8_R",isDat[["191031"]][,1]),"d13C"],"C4"))
foodDf = rbind(foodDf,cbind("Larvae on C4 food",isDat[["200420"]][grep("R12s_C4_d6_R",isDat[["200420"]][,1]),"d13C"],"C4"))
foodDf = rbind(foodDf,cbind("Larvae on C4 food",isDat[["200420"]][grep("R12s_C4_d7_R",isDat[["200420"]][,1]),"d13C"],"C4"))
foodDf = rbind(foodDf,cbind("Larvae on C4 food",isDat[["200420"]][grep("R12s_C4_d8_R",isDat[["200420"]][,1]),"d13C"],"C4"))
foodDf = rbind(foodDf,cbind("Larvae on C4 food",isDat[["200420"]][grep("R12s_C4_d9_R",isDat[["200420"]][,1]),"d13C"],"C4"))
foodDf = rbind(foodDf,cbind("Larvae on C4 food",isDat[["200420"]][grep("R12s_C4_d10_R",isDat[["200420"]][,1]),"d13C"],"C4"))
ggplot(foodDf,aes(x=factor(V1,unique(foodDf[,1])),y=as.numeric(V2),color=V3)) +
  geom_point(size=1,color="black") +
  geom_boxplot(alpha=0) +
  theme_minimal_grid() +
  scale_colour_manual(values=c("darkolivegreen4","deepskyblue2","darkgoldenrod2")) +
  ylim(c(-30, -10)) +
  xlab("") +
  ylab("d13C") +
  ggpubr::rotate_x_text()
#ggsave(paste0("D:/rad/publications/200602 isotope method/200608 food components.pdf"))


# R12s C3 and C4 all sample R vs TB and d13C vs ugC
rsDf = data.frame()
rsDf = rbind(rsDf,cbind("samp"="food","µgC"=0,"d13C"=isDat[["200420"]][grep("C3_food",isDat[["200420"]][,1]),"d13C"],"group"="Food C3"))
rsDf = rbind(rsDf,cbind("samp"="food","µgC"=0,"d13C"=isDat[["200420"]][grep("C4_food",isDat[["200420"]][,1]),"d13C"],"group"="Food C4"))
for (type in c("C3","C4")) {
  rsDat = isDat[["200420"]][grep("R12s",isDat[["200420"]][,1]),]
  sampSet = unique(unlist(strsplit(rsDat[,1]," "))[c(T,F)])
  sampSet = sampSet[grep(type,sampSet)]
  for (samp in sampSet) {
    if (length(grep("_TB",samp))>0) rsDf = rbind(rsDf,cbind("samp"=unlist(strsplit(gsub("_TB","",samp),"_"))[length(unlist(strsplit(gsub("_TB","",samp),"_")))],
                                                            "µgC"=rsDat[grep(samp,rsDat[,"ID"]),"µgC"],"d13C"=rsDat[grep(samp,rsDat[,"ID"]),"d13C"],"group"=paste("Cephalic complex",type,sep=" ")))
    if (length(grep("_R",samp))>0) rsDf = rbind(rsDf,cbind("samp"=unlist(strsplit(gsub("_R","",samp),"_"))[length(unlist(strsplit(gsub("_R","",samp),"_")))],
                                                           "µgC"=rsDat[grep(samp,rsDat[,"ID"]),"µgC"],"d13C"=rsDat[grep(samp,rsDat[,"ID"]),"d13C"],"group"=paste("Host tissues",type,sep=" ")))
  }
}
ggplot(rsDf,aes(x=factor(samp,level=unique(samp)),y=as.numeric(d13C),fill=group,color=group)) +
  geom_point(position = position_jitterdodge(jitter.width=0),color="black") +
  theme_minimal_grid() +
  xlab("") +
  geom_boxplot(alpha = 0) +
  ylab("d13C") +
  ylim(c(-30,-10)) +
  scale_color_manual(values=c("deepskyblue2","darkgoldenrod2","deepskyblue","darkgoldenrod","deepskyblue4","darkgoldenrod4"))
#ggsave(paste0("D:/rad/publications/200602 isotope method/200603 Food R12s d13C R and TB.pdf"))


# d6 - d8 masses and isotope measurements
d6d8df = data.frame()
d6d8df = rbind(d6d8df,cbind("gen"="R12s","tis"="Cephalic complex","samp"="day 0-6: C3",isDat[["191031"]][grep("R12s_C3_d6_TB",isDat[["191031"]][,1]),c("d13C","µgC")]))
d6d8df = rbind(d6d8df,cbind("gen"="R12s","tis"="Remaining host","samp"="day 0-6: C3",isDat[["191031"]][grep("R12s_C3_d6_R",isDat[["191031"]][,1]),c("d13C","µgC")]))
d6d8df = rbind(d6d8df,cbind("gen"="R12s","tis"="Cephalic complex","samp"="day 0-8: C3",isDat[["191031"]][grep("R12s_C3_d8_TB",isDat[["191031"]][,1]),c("d13C","µgC")]))
d6d8df = rbind(d6d8df,cbind("gen"="R12s","tis"="Remaining host","samp"="day 0-8: C3",isDat[["191031"]][grep("R12s_C3_d8_R",isDat[["191031"]][,1]),c("d13C","µgC")]))
d6d8df = rbind(d6d8df,cbind("gen"="R12s","tis"="Cephalic complex","samp"="day 0-6: C4",isDat[["191031"]][grep("R12s_C4_d6_TB",isDat[["191031"]][,1]),c("d13C","µgC")]))
d6d8df = rbind(d6d8df,cbind("gen"="R12s","tis"="Remaining host","samp"="day 0-6: C4",isDat[["191031"]][grep("R12s_C4_d6_R",isDat[["191031"]][,1]),c("d13C","µgC")]))
d6d8df = rbind(d6d8df,cbind("gen"="R12s","tis"="Cephalic complex","samp"="day 0-8: C4",isDat[["191031"]][grep("R12s_C4_d8_TB",isDat[["191031"]][,1]),c("d13C","µgC")]))
d6d8df = rbind(d6d8df,cbind("gen"="R12s","tis"="Remaining host","samp"="day 0-8: C4",isDat[["191031"]][grep("R12s_C4_d8_R",isDat[["191031"]][,1]),c("d13C","µgC")]))
d6d8df = rbind(d6d8df,cbind("gen"="R12s","tis"="Cephalic complex","samp"="day 0-6: C3, day 6-8: C4",isDat[["191031"]][grep("R12s_C3-C4_d6-d8_TB",isDat[["191031"]][,1]),c("d13C","µgC")]))
d6d8df = rbind(d6d8df,cbind("gen"="R12s","tis"="Remaining host","samp"="day 0-6: C3, day 6-8: C4",isDat[["191031"]][grep("R12s_C3-C4_d6-d8_R",isDat[["191031"]][,1]),c("d13C","µgC")]))
d6d8df = rbind(d6d8df,cbind("gen"="R12s","tis"="Cephalic complex","samp"="day 0-6: C3, day 6-8: C4 sugar only",isDat[["191031"]][grep("R12s_C3-C4sug_d6-d8_TB",isDat[["191031"]][,1]),c("d13C","µgC")]))
d6d8df = rbind(d6d8df,cbind("gen"="R12s","tis"="Remaining host","samp"="day 0-6: C3, day 6-8: C4 sugar only",isDat[["191031"]][grep("R12s_C3-C4sug_d6-d8_R",isDat[["191031"]][,1]),c("d13C","µgC")]))
d6d8df = rbind(d6d8df,cbind("gen"="R12s","tis"="Cephalic complex","samp"="day 0-6: C3, day 6-8: C4 yeast only",isDat[["191031"]][grep("R12s_C3-C4yea_d6-d8_TB",isDat[["191031"]][,1]),c("d13C","µgC")]))
d6d8df = rbind(d6d8df,cbind("gen"="R12s","tis"="Remaining host","samp"="day 0-6: C3, day 6-8: C4 yeast only",isDat[["191031"]][grep("R12s_C3-C4yea_d6-d8_R",isDat[["191031"]][,1]),c("d13C","µgC")]))
d6d8df = rbind(d6d8df,cbind("gen"="R12s","tis"="Cephalic complex","samp"="day 0-6: C3, day 6-8: starved",isDat[["191031"]][grep("R12s_C3-mq_d6-d8_TB",isDat[["191031"]][,1]),c("d13C","µgC")]))
d6d8df = rbind(d6d8df,cbind("gen"="R12s","tis"="Remaining host","samp"="day 0-6: C3, day 6-8: starved",isDat[["191031"]][grep("R12s_C3-mq_d6-d8_R",isDat[["191031"]][,1]),c("d13C","µgC")]))
datPlot = d6d8df[d6d8df[,"gen"]=="R12s",]
ggplot(datPlot,aes(x=factor(samp,level=unique(samp)),y=as.numeric(d13C),fill=tis,color=tis)) +
  geom_point(position = position_jitterdodge(jitter.width=0),color="black") +
  theme_minimal_grid() +
  xlab("") +
  geom_boxplot(alpha = 0) +
  ylab("d13C") +
  ylim(c(-30,-10)) +
  labs(subtitle=paste("RasV12,scribble clones in eye-antennal disc")) +
  ggpubr::rotate_x_text()
#ggsave(paste0("D:/rad/publications/200602 isotope method/200610 R12s d6d8 d13c all.pdf"))
datPlot = d6d8df[d6d8df[,"gen"]=="R12s" & d6d8df[,"tis"]=="Cephalic complex",]
ggplot(datPlot,aes(x=factor(samp,level=unique(samp)),y=as.numeric(µgC))) +
  geom_point(color="black") +
  theme_minimal_grid() +
  xlab("") +
  geom_boxplot(alpha = 0) +
  ylab("Cephalic complex, µgC") +
  ylim(c(0,50)) +
  labs(subtitle=paste("RasV12,scribble clones in eye-antennal disc")) +
  geom_signif(comparisons=as.list(as.data.frame(combn(unique(datPlot[,3]),2))),test="t.test",
              step_increase=0.025,tip_length=0,size=0.3,textsize=2,vjust=1.4,color="black") +
  ggpubr::rotate_x_text()
#ggsave(paste0("D:/rad/publications/200602 isotope method/200610 R12s d6d8 ugc TB.pdf"))


# Calculating contribution to growth, d6-d8
plotDf = data.frame()
tmpDat = isDat[["191031"]][grep("R12s",isDat[["191031"]][,1]),]
selSets = c("C3-C4_d6-d8_TB","C3-C4sug_d6-d8_TB","C3-C4yea_d6-d8_TB","C3-mq_d6-d8_TB")
ugcPrev = mean(tmpDat[grep("_d6_TB",tmpDat[,"ID"]),"µgC"])
ugcNext = mean(tmpDat[grep("_d8_TB",tmpDat[,"ID"]),"µgC"])
ugcChg = ugcNext - ugcPrev
c3prev = mean(tmpDat[grep("C3_d6_TB",tmpDat[,"ID"]),"d13C"])
c3next = mean(tmpDat[grep("C3_d8_TB",tmpDat[,"ID"]),"d13C"])
c4next = mean(tmpDat[grep("C4_d8_TB",tmpDat[,"ID"]),"d13C"])
for (selSet in selSets) {
  for (selrow in grep(selSet,tmpDat[,"ID"])) {
    ugSamp = as.numeric(tmpDat[selrow,"µgC"])
    ugcPrev = ugSamp - ugcChg
    ugAdded = ugSamp - ugcPrev
    ifAllTissues = ((ugcPrev*c3prev)+(ugAdded*c3next))/(ugcPrev+ugAdded)
    ifAllFood = ((ugcPrev*c3prev)+(ugAdded*c4next))/(ugcPrev+ugAdded)
    d13sel = as.numeric(tmpDat[selrow,"d13C"])
    ratioFood = (ifAllTissues - d13sel) / (ifAllTissues-ifAllFood)
    ratioTissues = 1-ratioFood
    plotDf = rbind(plotDf,cbind("samp"=paste0("R12s\n",paste(strsplit(selSet,"_")[[1]],collapse="\n")),
                                "ratio"="Food","val"=ratioFood,"ugCadded"=ratioFood*ugAdded))
    plotDf = rbind(plotDf,cbind("samp"=paste0("R12s\n",paste(strsplit(selSet,"_")[[1]],collapse="\n")),
                                "ratio"="Host","val"=ratioTissues,"ugCadded"=ratioTissues*ugAdded))
  }
}
ggplot(plotDf,aes(x=factor(samp,unique(plotDf[,"samp"])),y=as.numeric(ugCadded),color=ratio)) +
  geom_boxplot(alpha=1) +
  geom_point(position = position_jitterdodge(jitter.width=0)) +
  theme_minimal_grid() +
  xlab("") + 
  ylab("µg carbon added to tumour from nutrient source")
#ggsave(paste0("D:/rad/publications/200602 isotope method/200604 d6_d8 ugC R12s.pdf"))
```
